# SARS-CoV-2 multi-antigen protein microarray for detailed characterization of antibody responses in COVID-19 patients

**DOI:** 10.1101/2022.10.14.512324

**Authors:** Alev Celikgil, Aldo B. Massimi, Antonio Nakouzi, Natalia G. Herrera, Nicholas C. Morano, James H. Lee, Hyun ah Yoon, Scott J. Garforth, Steven C. Almo

**Author notes:** Corresponding authors, (SCA), (AC).

## Abstract

Antibodies against severe acute respiratory syndrome coronavirus 2 (SARS-CoV-2) target multiple epitopes on different domains of the spike protein, and other SARS-CoV-2 proteins. We developed a SARS-CoV-2 multi-antigen protein microarray with the nucleocapsid, spike and its domains (S1, S2), and variants with single (D614G, E484K, N501Y) or double substitutions (N501Y/Deletion69/70), allowing a more detailed high-throughput analysis of the antibody repertoire following infection. The assay was demonstrated to be reliable and comparable to ELISA. We analyzed antibodies from 18 COVID-19 patients and 12 recovered convalescent donors. S IgG level was higher than N IgG in most of the COVID-19 patients, receptor-binding domain of S1 showed high reactivity, but no antibodies were detected against heptad repeat domain 2 of S2. Furthermore, antibodies were detected against S variants with single and double substitutions in COVID-19 patients who were infected with SARS-CoV-2 early in the pandemic. Here we demonstrated that SARS-CoV-2 multi-antigen protein microarray is a powerful tool for detailed characterization of antibody responses, with potential utility in understanding the disease progress and assessing current vaccines and therapies against evolving SARS-CoV-2.

## 1. Introduction

Severe acute respiratory syndrome coronavirus 2 (SARS-CoV-2) has caused a pandemic with significant global impact since it was first identified in December 2019. The immune responses of infected individuals have been studied to understand pathogenesis of asymptomatic to severe disease using serological assays, primarily ELISA. Initially, the major antigen target for these assays was nucleocapsid (N) [1–4], one of the antigenic structural proteins; later, the focus has been on the spike (S), another antigenic structural protein [5, 6].

S glycoprotein consists of S1 and S2 domains; the S1 domain, which mediates binding to the host receptor angiotensin-converting enzyme 2 and the S2 domain, responsible for the fusion of host and viral cell membrane using six-helical bundle structure formed with heptad repeat (HR) domains [7]. Antibodies target multiple epitopes on the S protein and yet only some of them can induce neutralizing response against SARS CoV-2 [8, 9]. Specifically, neutralization has been shown to be proportional to the titer of antibodies that target receptor-binding domain (RBD) of S protein [10].

As SARS-CoV-2 evolves, new variants are emerging with mutations that potentially affect antigenicity and pose new challenges for vaccines and therapies. These variants have multiple mutations in RBD and non-RBD subdomains including D614G, E484K, N501Y, Deletion69/70, located in S1 domain of S, and some variants confer resistance to therapeutic monoclonal antibodies or to the antibodies elicited by vaccination [11]. While D614G and N501Y mutations showed minimal impact on neutralization, E484K mutation reduced neutralization by the monoclonal antibodies, convalescent plasma (CP) therapy and post-vaccination sera [12–15]. Since immunogenic epitopes are distributed across the entire S protein, more detailed characterization of antibody signatures is crucial not only to understand the disease progress, but also to evaluate CP therapy and vaccine responses for protection against infection with S variants. For this purpose, we developed a SARS-CoV-2 multi-antigen protein array with structural proteins N, S and its domains (S1, S2, RBD, HR2). This assay allows parallel testing for antibody responses to different targets and domains of those targets using a very small sample volume. We were able to determine the antibody signatures of multiple COVID-19 patients, as well as CP donors who recovered from COVID-19. Furthermore, we evaluated the binding of polyclonal antibodies in CP samples from individuals who were infected with SARS-CoV-2 early in the pandemic to S protein containing the clinically relevant substitutions D614G, E484K, N501Y and N501Y/Deletion69/70.

## 2. Materials and methods

### 2.1. Study design

We generated a SARS-CoV-2 protein microarray with two most antigenic structural proteins N and S along with S domains (S1, S2), RBD subdomain of S1, HR2 subdomain of S2, S variants with single (D614G, E484K, N501Y) and double (N501Y/Deletion69/70) substitutions. After the quality of immobilized proteins were checked, the assay was validated with ELISA using eight CP donor plasma. Multiple samples were screened in parallel on a single microarray slide designed with 12-16 subarrays. We analyzed the antigen-specific IgG, IgM and IgA responses in eighteen COVID-19 patients. Eleven of these patients were also evaluated for the antibody response to S variants. In addition, four COVID-19 patients were assessed before and after the therapy along with their donors. (**S1 Fig**).

### 2.2. Plasmids

Ectodomain of SARS-CoV-2 S (residues 1-1208) in pCAGGS vector was kindly provided by McLellan and coworkers [16]. S2 domain (residues 687-1208) in pCAGGS vector was kindly provided by John Lai (Einstein). S1 domain (residues 13-685), RBD of S1 (residues 319-541) and HR2 of S2 (residues 1163-1202) were amplified by polymerase chain reaction (PCR) using SARS-CoV-2 S as template and subcloned into pcDNA3.3 vector with a C-terminal hexahistidine tag using In-Fusion cloning technology (Takara Bio USA, Inc.). RBD of SARS-CoV S was cloned into pcDNA3.3 vector from pcDNA3.1-SARS-Spike (Addgene plasmid#145031). SARS-CoV-2 N was cloned into a pSGC-his vector as described in our previous paper [17]. Single and double mutations were introduced using PCR and In-Fusion with the McLellan S expression construct.

### 2.3. Antigen expression and purification

S and N proteins were expressed and purified as previously described in Herrera et al. [17]. S1, S2, HR2 and S variants were expressed in ExpiCHO-S™ cells as described for full-length S. RBD of SARS-CoV2 and SARS-CoV S were transiently expressed in FreeStyle™ 293-F cells (Thermo Fisher Scientific). Cells were resuspended at 1×10^6^ per mL with fresh media on the day of transfection. DNA (0.5mg/ml) and Polyethylenimine (2mg/ml; PEI, Polysciences, Inc., 23966) were mixed in 1XPBS and incubated for 15 minutes at room temperature. The DNA/PEI mixture was added to the cells dropwise and cells were incubated in a shaker at 37°C and 5% CO2 [18]. 24 hours post-transfection, valproic acid salt (Sigma-Aldrich, P4543-100G) suspended in media (1/3 of starting volume) was added to the cells at final concentration of 3mM [19]. The cells were harvested on day 7 post-transfection. S domains and variants were purified similar to S protein using nickel resin (10 mL/L, His60 Ni2+ superflow resin, Takara cat# 635664) with wash buffer (25 mM MES, 150 mM NaCl, 10% glycerol, 50 mM Arg-Cl, 5mM imidazole, pH 6.5) and elution buffer (25 mM MES, 150 mM NaCl, 10% glycerol, 100 mM Arg-Cl, 0.5 M imidazole, pH 6.50). The eluates were concentrated using Amicon centrifugal units (EMD Millipore) and dialyzed against 50 mM Tris, 250 mM NaCl, pH 8.0 for 2 hours at room temperature and then overnight at 4°C. Proteins were analyzed by SDS-PAGE. Protein concentrations were determined using UV absorbance at 280nm; extinction coefficient was calculated from amino acid sequence using Expasy online ProtParam.

### 2.4. Sera/plasma samples

Remnant blood samples were available from patients who were positive by PCR for SARS-CoV-2 and received CPT as part of United States Expanded Access Program (EAP) [20] in April and May 2020 as described in Yoon et al. [21]. They were symptomatic for 3-7 days and hospitalized with severe COVID-19 for 3 days or less before the therapy. The remnant sera were obtained before and after transfusion (day 0, 1, 3, 7). CP donor samples were collected from individuals who were positive by PCR for SARS-CoV-2 and symptom free for at least 14 days before the plasma collection as part of an institutional donor program conducted in March-April 2020 as described previously [20] and used in EAP or CONTAIN COVID-19 trial [22]. Control samples were obtained from healthy individuals at New York Blood Center (NYBC) between December 2019 and February 2020. Samples were heat inactivated at 56°C for 30 minutes and stored at 4°C for short term or −80°C for long term.

The retrospective cohort study, the donor plasma procurement protocol, and the use of the EAP were approved by the Albert Einstein College of Medicine (AECOM) IRB. The retrospective cohort study that included collection of remnant blood samples of patients was approved by the AECOM IRB for human subjects with a waiver of informed consent.

### 2.5. Multi-antigen Protein Array Production and Processing

A protein array of SARS-CoV-2 antigens was generated with full-length N and S, S1, S2, RBD and HR2 domains of S protein. Negative controls included RBD of SARS-CoV S, 1XPBS, sample buffer and human acetylcholinesterase. Human immunoglobulin (Ig) isotype controls IgG, IgM and IgA were printed as reference to normalization for each sample, and to confirm detection by the secondary antibodies.

Each slide was printed with twelve identical subarrays, and used for serum dilutions or multiple serum screening. Target antigens were adjusted to between 0.3-7.2 femtomol (fmol), and Ig isotype controls 0.15-3.6 fmol per spot. Microarray slides were generated with aminosilane-coated slides using a piezoelectric printer (Arrayjet, Edinburgh, UK) and processed as described previously [17]. To determine levels of immobilized proteins, one slide was probed with Alexa Fluor® 647 anti-his antibody (1:250, BioLegend, 362611). Sera/plasma samples were single or serial diluted (three-fold dilutions from 1:90) in 250 μl 5% milk, PBS, 0.2% Tween-20.

### 2.6. Data analysis

Data was processed using GenePix® Pro7 software (Molecular Devices). Each sample was corrected for background by subtracting the raw spot intensity of negative control protein from every sample spot. The results were normalized relative to corresponding concentration of Ig isotype controls in each subarray to minimize the variations that occur during sera and array processing. The mean fluorescence intensity ± standard deviation (SD) was calculated from the replicates for each antigen concentration.

Results were plotted using GraphPad Prism software version 8.4.3. Nonlinear regression model was used for dilution-response curves of serum samples. The bar plots and heatmap were used to display antibody levels of multiple samples against antigen targets, CPT time points or Ig isotypes. The data was also represented as the ratio of antigen specific antibody levels.

### 2.7. ELISA

SARS-CoV-2 N, S2, RBD, and HR2 protein-binding IgG were measured by ELISA using donor CP as previously described in Bortz et al [23].

## 3. Results

### 3.1. Generation and validation of SARS-CoV-2 multi-antigen protein microarray

We initially generated a protein array of SARS-CoV-2 antigens with structural proteins N, S, S domains (S1, S2) and subdomains (RBD, HR2). Later, we extended the array to S variants with single (D614G, E484K, N501Y) and double mutations

(N501Y/Deletion69/70). While N protein was produced from *E. coli* cells, we used mammalian cells to express S protein, its domains and variants with post-translational modifications (PTM). Representative SDS-PAGE analysis of proteins is shown in **Fig 1A**.

**Fig 1.**
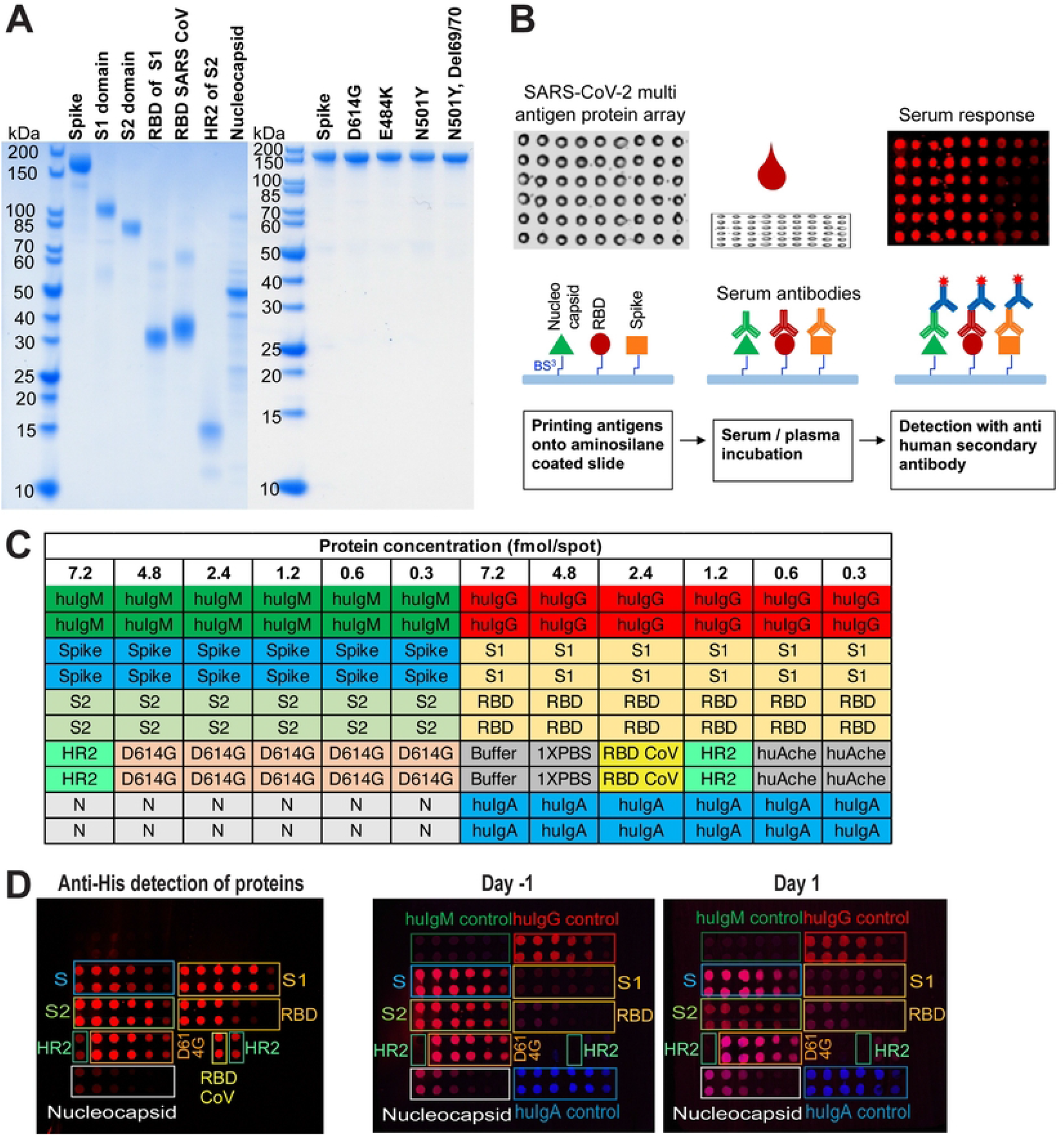
SARS-CoV-2 Multi-antigen protein array production and processing. (A) Purified array proteins analyzed by SDS-PAGE under reducing conditions. (B) Schematics of workflow. (C) A representative protein array template showing SARS-CoV-2 antigen target proteins, human immunoglobulin controls (huIgM, huIgG, huIgA), buffer and negative controls along with the protein amounts and duplicates of each sample. (D) Anti-his staining of antigen target proteins immobilized on the microarray slide, and serum antibody detection of a COVID-19 patient before (Day −1) and after (Day 1) convalescent plasma transfusion using secondary antibodies for IgG (red) and IgA (blue) (two-color image). huIg control proteins do not have his tags; N, nucleocapsid; S, spike protein.

After a concentration series of antigens, Ig isotype controls (IgG, IgM and IgA) and negative controls were arrayed onto the microarray slide (**Fig 1B and 1C**), the levels of immobilized proteins were determined using antibodies against the his tag (**Fig 1D**). A pattern of decreasing signal intensities correlated to antigen amount was observed, and no signal was detected from buffer spots. Multiple samples were screened in parallel on a single microarray slide designed with twelve subarrays. To visualize and analyze the human antibodies bound to the antigens on the slide, fluorescent-labeled secondary anti-human antibodies were used. Representative images are shown as two-color images (IgG/IgA) in **Fig 1D**.

### 3.2. Assay development

To identify a serum dilution that gives reliable detection for multiple samples with different antibody titers, 11 dilutions of a single sample were tested for IgG response to S and N proteins (**Fig 2A and B**). Next, 21 samples were assayed at the 3 highest dilutions (1/90, 1/270, 1/810) (**Fig 2C**). At all three dilutions, antibody response to S and N proteins could be detected in the sera/plasma; antibodies were not detected in samples from control subjects. 1/270 dilution was chosen for further analysis of multiple samples to avoid the false negative results observed with low titer samples at higher dilutions.

**Fig 2.**
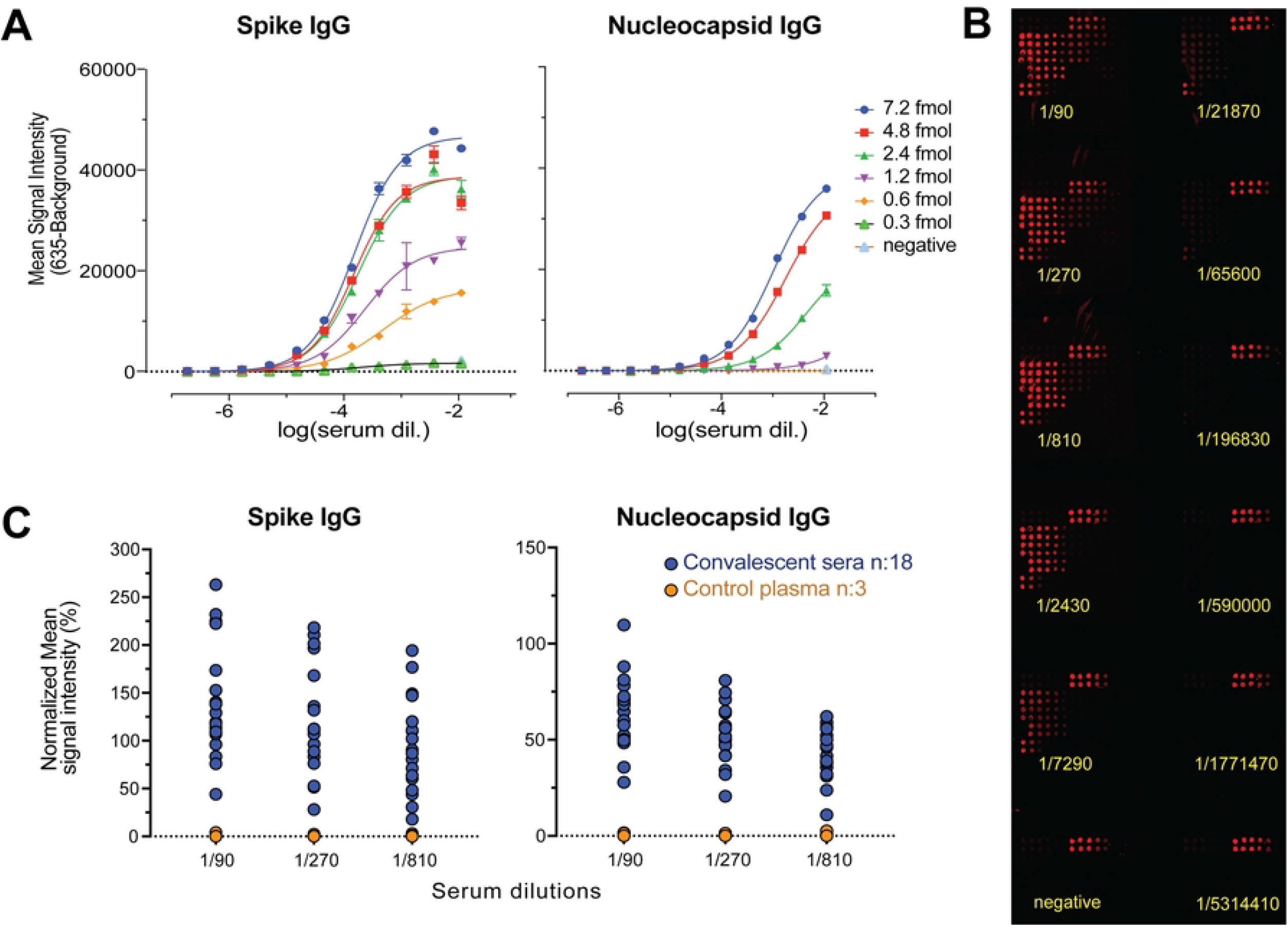
Serial dilution test to identify a single dilution that gives reliable detection for multiple sera/plasma samples. (A) Spike and nucleocapsid IgG levels are shown for single serum sample at 11 serial dilutions against six protein concentrations. Mean values ± SD are shown for duplicates. (B) Representative image of protein microarray printed with 12 identical subarrays that was used for IgG detection in a single sample with three-fold serial dilutions. (C) Spike and nucleocapsid IgG levels are shown for multiple samples (n:21) with three highest serial dilutions. The results are normalized relative to corresponding concentration of IgG isotype controls and mean values are calculated for the replicates of target antigen for each sample.

### 3.3. Protein array and ELISA comparison

We also compared our assay to ELISA which requires multiple plates to screen samples against different targets. When the same target antigens and same donor plasma samples (CONTAIN, **Table 1**) were used, results from protein microarray assay with 1/270 serum dilution and 2.4 fmol of each target antigen were comparable to ELISA which uses antigen concentration at saturation level (**S2 Fig**). Both assays identified the same samples with high levels of IgG for N, S2 and RBD, and also that they were seronegative for HR2 subdomain of S2.

**Table 1.** Information on sera/plasma samples tested in this study.

| Study group | *n* | Patient status | Characteristics | Source | Sample collection data |
| --- | --- | --- | --- | --- | --- |
| Patient | 18 | Severe and/or life threatening COVID-19 | Symptomatic for 3-7 days prior to transfusion | EAP_MMC<br>Yoon et al.<br>2021 | April - May<br>2020 |
| Donor | 8 | Recovered from COVID-19 | Asymptomatic for at least 14 days prior to sample collection | CONTAIN<br>Ortigoza et al. 2022 | March - April<br>2020 |
| Donor | 4 | Recovered from COVID-19 | Asymptomatic for at least 14 days prior to sample collection | EAP_MMC<br>Yoon et al.<br>2021 | March - April<br>2020 |
| Control | 3 | Not tested for SARS-CoV-2 | Pre-pandemic | NYBC | December 2019<br>- February 2020 |

Abbreviations: COVID-19, coronavirus disease 2019; EAP, expanded access protocol; MMC, Montefiore Medical Center; NYBC, New York Blood Center.

### 3.4. Antibody signatures of COVID-19 patients

Antibody responses of eighteen COVID-19 patients who were infected with SARS-CoV-2 between April and May 2020 were analyzed, in parallel with three control plasma samples of unknown infection history that were obtained between December 2019 and February 2020 (**Table 1**). Multi-antigen screening showed higher IgG for S than N in 14 of the COVID-19 patients (**Fig 3**). Qualitative analysis of antibody responses to S domains (S1, S2) and subdomains (RBD, HR2) showed higher S2 IgG than S1 and RBD IgG in most of the samples (16 and 11 samples, respectively). IgM and IgA reactivity in the patient sera were overall less than IgG (**S3A Fig**). S1/RBD IgG ratio was low in all of the samples, showing antibody responses to S1 domain was mostly targeting RBD (**S3B Fig**). We observed a wide range of S/RBD and S2/RBD ratios between individuals indicating different levels of serum antibodies targeting non-RBD regions on S2 compare to RBD.

**Fig 3.**
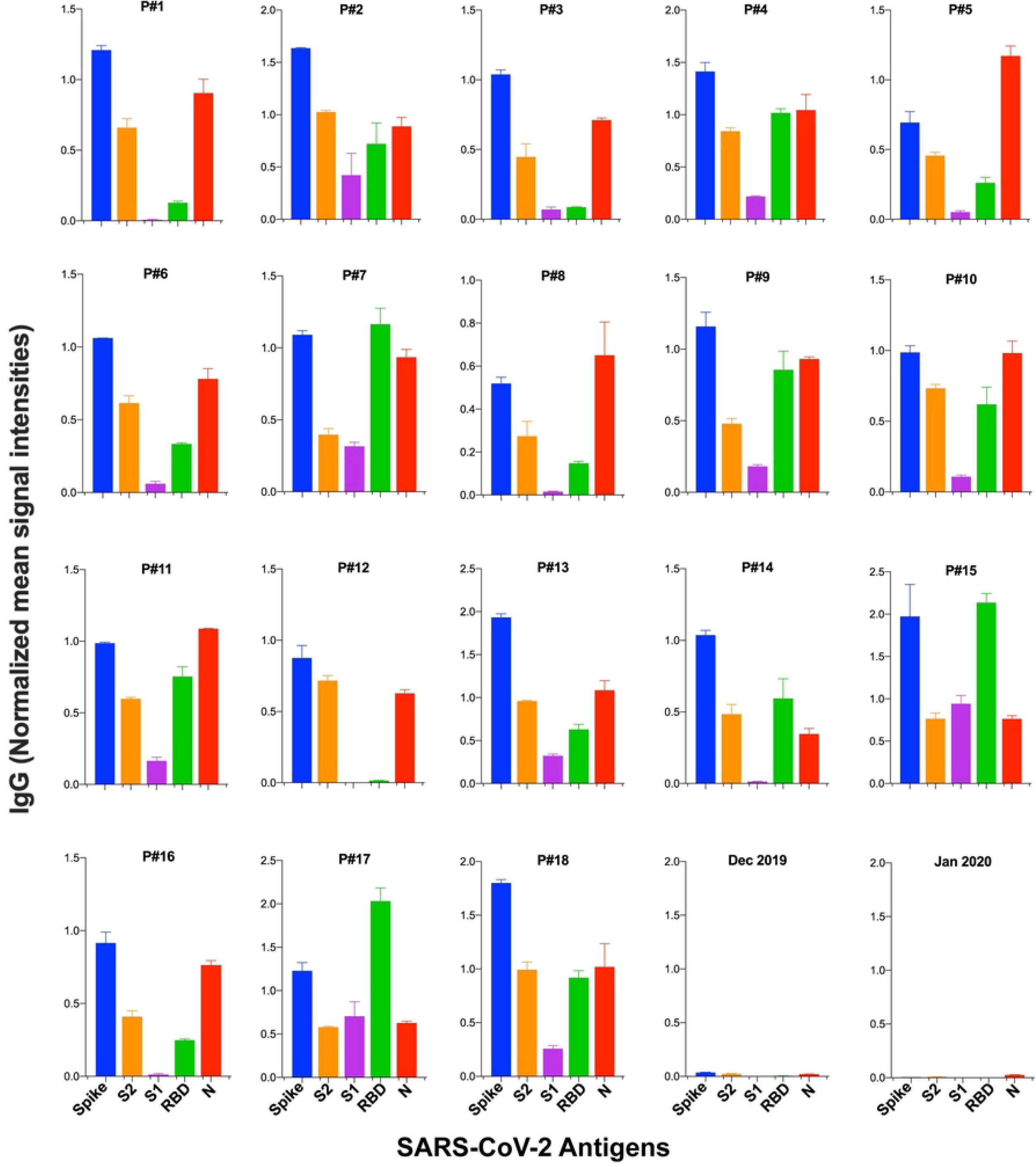
Serum IgG levels against SARS-CoV-2 antigens in COVID-19 patients. Serum IgG levels are shown against spike, its domains (S1, S2, RBD) and nucleocapsid (N) for 18 patients sera (P#1-18) and 2 control plasma samples. Signal intensities are normalized to human IgG isotype control for each sera/plasma and mean values ± SD are calculated for duplicates.

### 3.5. Antibody signatures of CPT donors and recipients

Antibody signature of CP has been proposed to affect the efficacy of CPT [24, 25]. We screened samples from four COVID-19 patients and their donors who recovered from COVID-19 (EAP_MMC, **Table 1**) against SARS-CoV-2 S, S2, RBD, HR2 and N before and after CP transfusion. Two of the CPT recipients died 3 days (patient#1) and 18 days (patient#2) post-transfusion; two recovered 52 days (patient#3) and 2 days (patient#4) post-transfusion. While we observed differences in antigen specific antibody responses between patients and donors, we were also able to monitor the changes in antibody levels at different time points post-CPT (**S4 Fig**). Although all the patient samples were seropositive for S2 IgG, we did not observe antibody response to HR2 subdomain of S2 in any of the samples.

### 3.6. Antibody responses to spike variants

In order to begin to address whether patient’s antibodies against SARS-CoV-2 from early pandemic could show response to later isolates, we extended our protein microarray panel to S variants with single mutations D614G (in CTD), E484K (in RBD), N501Y (in RBD) and double mutations N501Y/Del69/70 (in RBD/NTD) in S1 domain (**S5 Fig**). We screened eleven COVID-19 patients along with two control samples (EAP_MMC and NYBC, **Table 1**). While we observed serum responses to S variants with single and double mutations (**Figs 4A and 4B**), the IgG bound to D614G variant was ~2-fold lower than wildtype S-specific IgG in all patient samples tested (**Fig 4C**). IgG bound to E484K, N501Y, and N501Y/Del69-70 variants were comparable to wildtype S-specific IgG levels in most of the samples. Only one patient had slightly higher antibody response to these three variants than wildtype S (patient#2). In three patients (P#7, P#9, P#11), IgG response to E484K, which is in RBD domain, was less than the response to the other RBD mutation N501Y (**Fig 4B**). IgM levels were uniformly lower than IgG, which could be due to the time interval between symptom onset and the sample being taken. Although the patients 1, 6, 8 and 11 had higher levels of S-specific IgM compared to other patients, they were still less than IgG. In these samples, the differences between E484K and N501Y or N501Y/Del69/70 were higher for IgM than IgG, however the antibody levels were low.

**Fig 4.**
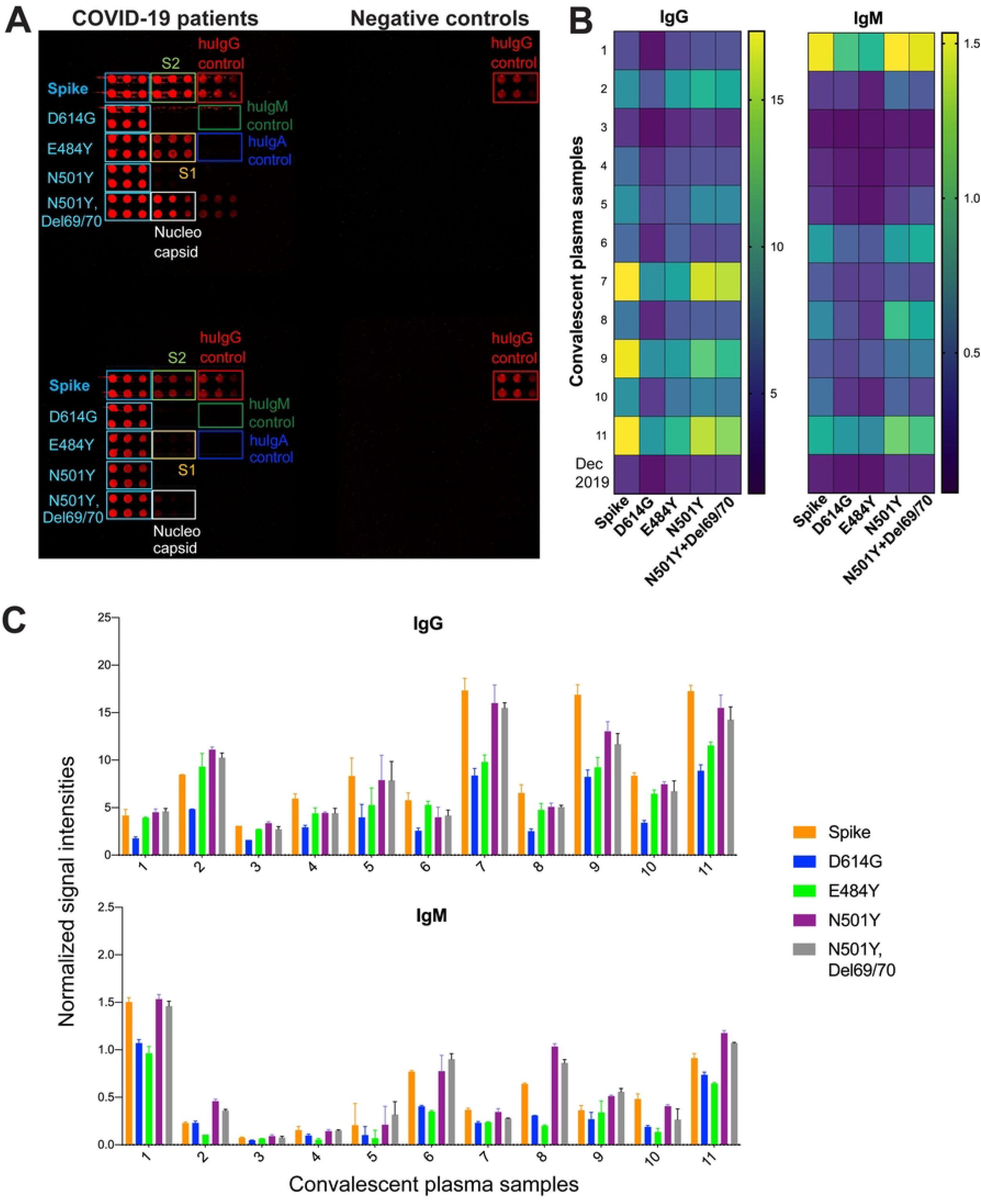
Antibody responses to S variants with single and double mutations in sera from COVID-19 patients from the beginning of the pandemic. (A) Representative image of four subarrays from SARS-CoV-2 protein microarray slide used for screening of two COVID-19 patients’ sera and two control plasma samples. Proteins are printed as 4.8 - 2.4 - 1.5 fmol in duplicates. IgG (red) responses are shown against S variants with single mutation D614G, E484K, N501Y and double mutations N501Y with Del69/70 along with S, S1, S2 and N. (B) Heatmap showing IgG and IgM profiles of patients’ sera (n:11) and control plasma (Dec 2019) against 2.4 fmol of each S variant. Signal intensities from duplicates are averaged. (C) Antibody levels are shown as bar plots and data are represented as normalized mean signal intensities ± SD.

## 4. Discussion

For detailed characterization of antibody responses against SARS-CoV-2 antigens, we developed a multi-antigen protein array initially with the two most immunogenic SARS-CoV-2 antigens S and N. Since antibodies target multiple epitopes on the S protein, and the antigenicity of these epitopes change due to mutagenesis in emerging SARS-CoV-2 variants [11], we extended the array to S domains and variants with single or double substitutions. S protein is heavily glycosylated, and its expression in non-mammalian systems might fail the detection of antibodies to glycosylated determinants, therefore we produced the proteins from mammalian cells for protein microarray fabrication [16, 26]. We also demonstrated that protein microarrays are comparable to ELISAs, which have been used widely for serological testing during the pandemic. Both assays not only indicated that same samples are seropositive or seronegative, but also showed similarity for the sample groups with high-medium-low levels of antibodies. The difference in response seen between the protein array and the ELISA is simply because of the high, fixed concentration of immobilized protein utilized in the ELISA. This could increase sensitivity of the ELISA relative to the protein array, particularly for low affinity antibodies, but may reduce the linearity of the response when comparing antigens that are at opposite ends of the detection spectrum.

In our serum screening, all of the individuals who were infected with SARS-CoV-2 had reactivity with full-length S protein, but binding to distinct S domains differed. Furthermore, we showed that sera from early in the pandemic contained antibodies capable of binding SARS-CoV-2 S protein containing the clinically relevant substitutions D614G, E484K, N501Y and N501Y/Deletion69/70, prevalent in clinical isolates from later in the outbreak.

In this study, our screening of COVID-19 patients showed high S-specific IgG levels compare to N-specific IgG in most of the patients as shown in previous studies [25, 27]. We also studied the immunogenic features of specific subdomains; RBD in S1 domain and HR2 in S2 domain of S. RBD subdomain of S1 is a highly variable domain of S protein. Low ratio of S1 to RBD IgG observed in patients sera was suggesting that RBD is the antigenic determinant of S1. HR2 subdomain of S2 plays a major role in the viral fusion to the cell membrane, and is a target of fusion inhibitors against SARS-CoV-2 [28]. We did not observe any signal against HR2 in samples, even those with high levels of S2-specific antibodies. The wide range of S/RBD and S2/RBD IgG ratios in samples suggest a different distribution of dominant epitopes in S domains between individuals.

Although RBD is the main antigenic region in S1 domain, most of the COVID-19 patients showed higher levels of antibody binding with non-RBD targets in S2 domain, as reported in data from other studies [8, 29], which showed similar signature as fulllength S protein between patients.

IgM and IgA antibodies were low in most of the patient samples; presence or absence may reflect differences in time of exposure to the virus and individual’s immune responses. While presence of IgM is indication of early disease, IgA is detected in more severe disease outcomes [30–32]. Target and isotype differences of serum antibodies combined with clinical features may be useful predictors of disease progress for individuals infected with SARS-CoV-2. Furthermore, immunoprofiling using multi-antigen protein arrays with only couple microliter of sera could help to detect distinct antibody characteristics of CPT donors and recipients, and monitor the disease progress post-therapy. An advantage of the protein chip for detecting antibodies is the ability to assay many samples, in this case time points, in parallel. Whilst it is clear that the patients tested showed different levels of antibodies against the antigens tested (Fig. 5), the sample size would have to be greatly expanded to detect a possible correlation with CP therapy.

Although SARS-CoV-2 protein microarrays were used for serum screening of CP in previous studies [33, 34], S variants were not included in the arrays, and S2 was not expressed in mammalian systems which could affect antigenicity due to missing PTMs. As new SARS-CoV-2 variants are emerging, reinfection cases are increasing all around the world. These variants have several mutations located in the subdomains of S protein (**S5 Fig**). From a few cases studied, Tillett et al. reported an asymptomatic reinfection case which was infected with two SARS-CoV-2 variants that both had single D614G substitution in S and several different mutations outside of S protein [35]. Prado-Vivar et al. reported a reinfection case with worse disease than the first infection [36]. This individual was infected with S variant with single D614G substitution at first infection, and wild type S at second infection. These individuals were infected with variants that had also several different mutations outside of S protein. There was no data on individuals’ antibody responses to S and its domains. In our screening of individuals infected with SARS-CoV-2 early in the pandemic, their response was mounted against the virus they were infected with, and their antibodies bound to other variants with single mutations D614G, E484K, N501Y and double mutations N501Y/Del69/70 in S1 domain of S. This is likely due to polyclonality of antibodies in the sera/plasma, targeting a broad range of epitopes, as previously suggested [14].

However, specific combinations of mutations in S protein, as well as in other immunogenic viral proteins could diminish the protective immunity from previous infection and increase the probability of reinfection. SARS-CoV-2 Beta (B.1.351) variant, which includes E484K, D614G, N501Y and other mutations in S, exhibited a reduced neutralization by plasma from individuals infected with SARS-CoV-2 early in the pandemic or vaccinated [11]. We observed some reduction in response to S with single D614G substitution in all of the samples, but only in a few samples to E484K, presumably reflects the differences between individuals’ antibody repertoire.

The strength of our study was screening each individual against multiple targets and concentrations in the same subarray with single small sample volume unlike ELISAs which requires multiple plates and sample aliquotes. Moreover, this assay could be used in a point-of-care setting to provide individuals’ full antibody signatures by targeting multiple antigens unlike most of the current point-of-care tests which targets single antigen.

Most importantly, our protein microarray, allowing parallel analysis of multiple variant proteins and their subdomains, is ideally suited for the analysis of the potential of patient antibodies to neutralize variants other than that which was responsible for the initial infection. Further, it is hoped that by extending the scope of the study and analyzing full spectrum of antibody responses, we can determine immunogenic antigens that are less prone to loss through viral evolution.

## Acknowledgements

We thank Liise-anne Pirofski for her insightful discussions and editorial input for the manuscript. We also thank Jason S. Mclellan for providing spike plasmid, and Jonathan R. Lai for sharing S2 plasmid.

## Supporting information

**S1 Fig. Study design for SARS-CoV-2 multi-antigen protein array.**

**S2 Fig. IgG responses to nucleocapsid protein (N) and S domains detected with protein microarray and ELISA.** Sera with highest antibody responses are highlighted with blue frame for each target antigen for both assays. (A) IgG responses to N, S2 domain, RBD and HR2 subdomains determined from protein array for eight convalescent sera along with negative and positive controls. Signal intensities from duplicates of each target antigen are averaged and normalized to corresponding concentration of human IgG isotype control. (B) IgG responses to N, S2, RBD and HR2 in the same eight sera are detected with ELISA in eleven three-fold serial dilutions starting from 1/100. Absorbance at 450nm is averaged for duplicates and represented using non-linear regression model.

**S3 Fig. Immunoprofiling COVID-19 patients against multiple SARS-CoV-2 antigens.** (A) Bar plots are showing IgG, IgM, and IgA response to 4.8 fmol of each S, S2, S1, RBD and N antigen for 18 patients who were infected early in the pandemic.

The signal intensities are normalized relative to corresponding concentration of Ig isotype controls and mean values are calculated for the replicates of target antigen for each plasma sample. (B) The ratios of S, S2 and S1 to RBD domain are shown as bar plots for IgG.

**S4 Fig. IgG profiles of four CP therapy donors and four recipients against SARS-CoV-2 proteins before and after CP therapy.** IgG serum responses are shown as percentages of normalized mean signal intensities against 4.8 fmol antigen proteins spike, S2, RBD, HR2 and nucleocapsid. CPT time points for the recipients are shown as one day before the transfusion, one and three days after the transfusion along with corresponding donor’s antibody levels (CP).

**S5 Fig. Shared Spike mutations in SARS-CoV-2 variants.** S variants’ shared mutations E484K, N501Y, D614G and Deletion69/70 are shown with their location on S1 domain of S protein. NTD, N-terminal domain; RBD, Receptor binding domain; CTD, C-terminal domain; FP, Fusion peptide; HR1, Heptad repeat 1; HR2, Heptad repeat 2.

## References

1. Guo L, Ren L, Yang S, Xiao M, Chang D, Yang F, et al. Profiling Early Humoral Response to Diagnose Novel Coronavirus Disease (COVID-19). Clin Infect Dis. 2020;71(15):778–85.

2. Gao T, Hu M, Zhang X, Li H, Zhu L, Liu H, et al. Highly pathogenic coronavirus N protein aggravates lung injury by MASP-2-mediated complement over-activation. medRxiv. 2020:2020.03.29.20041962.

3. Xiang F, Wang X, He X, Peng Z, Yang B, Zhang J, et al. Antibody Detection and Dynamic Characteristics in Patients With Coronavirus Disease 2019. Clinical infectious diseases: an official publication of the Infectious Diseases Society of America. 2020;71(8):1930–4.

4. Grant BD, Anderson CE, Williford JR, Alonzo LF, Glukhova VA, Boyle DS, et al. SARS-CoV-2 Coronavirus Nucleocapsid Antigen-Detecting Half-Strip Lateral Flow Assay Toward the Development of Point of Care Tests Using Commercially Available Reagents. Analytical Chemistry. 2020;92(16):11305–9.

5. Wajnberg A, Amanat F, Firpo A, Altman DR, Bailey MJ, Mansour M, et al. Robust neutralizing antibodies to SARS-CoV-2 infection persist for months. Science. 2020;370(6521):1227–30.

6. Buchholz UJ, Bukreyev A, Yang L, Lamirande EW, Murphy BR, Subbarao K, et al. Contributions of the structural proteins of severe acute respiratory syndrome coronavirus to protective immunity. Proc Natl Acad Sci U S A. 2004;101(26):9804–9.

7. Huang Y, Yang C, Xu XF, Xu W, Liu SW. Structural and functional properties of SARS-CoV-2 spike protein: potential antivirus drug development for COVID-19. Acta Pharmacol Sin. 2020;41(9):1141–9.

8. Lv H, Wu NC, Tsang OT, Yuan M, Perera R, Leung WS, et al. Cross-reactive Antibody Response between SARS-CoV-2 and SARS-CoV Infections. Cell Rep. 2020;31(9):107725.

9. Yarmarkovich M, Warrington JM, Farrel A, Maris JM. Identification of SARS-CoV-2 Vaccine Epitopes Predicted to Induce Long-Term Population-Scale Immunity. Cell Reports Medicine. 2020;1(3):100036.

10. Suthar MS, Zimmerman MG, Kauffman RC, Mantus G, Linderman SL, Hudson WH, et al. Rapid Generation of Neutralizing Antibody Responses in COVID-19 Patients. Cell Rep Med. 2020;1(3):100040.

11. Wang P, Nair MS, Liu L, Iketani S, Luo Y, Guo Y, et al. Antibody resistance of SARS-CoV-2 variants B.1.351 and B.1.1.7. Nature. 2021;593(7857):130–5.

12. Weisblum Y, Schmidt F, Zhang F, DaSilva J, Poston D, Lorenzi JC, et al. Escape from neutralizing antibodies by SARS-CoV-2 spike protein variants. Elife. 2020;9.

13. Wibmer CK, Ayres F, Hermanus T, Madzivhandila M, Kgagudi P, Oosthuysen B, et al. SARS-CoV-2 501Y.V2 escapes neutralization by South African COVID-19 donor plasma. Nat Med. 2021;27(4):622–5.

14. Rees-Spear C, Muir L, Griffith SA, Heaney J, Aldon Y, Snitselaar JL, et al. The effect of spike mutations on SARS-CoV-2 neutralization. Cell Rep. 2021;34(12):108890.

15. Zhou H, Dcosta BM, Samanovic MI, Mulligan MJ, Landau NR, Tada T. B.1.526 SARS-CoV-2 variants identified in New York City are neutralized by vaccine-elicited and therapeutic monoclonal antibodies. bioRxiv. 2021.

16. Wrapp D, Wang N, Corbett KS, Goldsmith JA, Hsieh CL, Abiona O, et al. Cryo-EM structure of the 2019-nCoV spike in the prefusion conformation. Science. 2020;367(6483):1260–3.

17. Herrera NG, Morano NC, Celikgil A, Georgiev GI, Malonis RJ, Lee JH, et al. Characterization of the SARS-CoV-2 S Protein: Biophysical, Biochemical, Structural, and Antigenic Analysis. ACS Omega. 2020;6(1):85–102.

18. Tom R, Bisson L, Durocher Y. Transient Expression in HEK293-EBNA1 Cells. In: Dyson M, Durocher Y, editors. In Methods Express: Expression Systems. 1st ed: Scion Publishing Ltd; 2007.

19. Backliwal G, Hildinger M, Kuettel I, Delegrange F, Hacker DL, Wurm FM. Valproic acid: a viable alternative to sodium butyrate for enhancing protein expression in mammalian cell cultures. Biotechnol Bioeng. 2008;101(1):182–9.

20. Joyner MJ, Carter RE, Senefeld JW, Klassen SA, Mills JR, Johnson PW, et al. Convalescent Plasma Antibody Levels and the Risk of Death from Covid-19. N Engl J Med. 2021;384(11):1015–27.

21. Yoon HA, Bartash R, Gendlina I, Rivera J, Nakouzi A, Bortz RH, 3rd, et al. Treatment of severe COVID-19 with convalescent plasma in Bronx, NYC. JCI Insight. 2021;6(4).

22. Ortigoza MB, Yoon H, Goldfeld KS, Troxel AB, Daily JP, Wu Y, et al. Efficacy and Safety of COVID-19 Convalescent Plasma in Hospitalized Patients: A Randomized Clinical Trial. JAMA Internal Medicine. 2022;182(2):115–26.

23. Bortz Robert H, Florez C, Laudermilch E, Wirchnianski Ariel S, Lasso G, Malonis Ryan J, et al. Single-Dilution COVID-19 Antibody Test with Qualitative and Quantitative Readouts. mSphere.6(2):e00224–21.

24. Herman JD, Wang C, Loos C, Yoon H, Rivera J, Eugenia Dieterle M, et al. Functional convalescent plasma antibodies and pre-infusion titers shape the early severe COVID-19 immune response. Nat Commun. 2021;12(1):6853.

25. Atyeo C, Fischinger S, Zohar T, Slein MD, Burke J, Loos C, et al. Distinct Early Serological Signatures Track with SARS-CoV-2 Survival. Immunity. 2020;53(3):524–32 e4.

26. Sanda M, Morrison L, Goldman R. N- and O-Glycosylation of the SARS-CoV-2 Spike Protein. Analytical Chemistry. 2021;93(4):2003–9.

27. Liu W, Liu L, Kou G, Zheng Y, Ding Y, Ni W, et al. Evaluation of Nucleocapsid and Spike Protein-Based Enzyme-Linked Immunosorbent Assays for Detecting Antibodies against SARS-CoV-2. J Clin Microbiol. 2020;58(6).

28. Zhu Y, Yu D, Yan H, Chong H, He Y. Design of Potent Membrane Fusion Inhibitors against SARS-CoV-2, an Emerging Coronavirus with High Fusogenic Activity. J Virol. 2020;94(14).

29. Rogers TF, Zhao F, Huang D, Beutler N, Burns A, He W-T, et al. Isolation of potent SARS-CoV-2 neutralizing antibodies and protection from disease in a small animal model. Science (New York, NY). 2020;369(6506):956–63.

30. Staats LAN, Pfeiffer H, Knopf J, Lindemann A, Fürst J, Kremer AE, et al. IgA2 Antibodies against SARS-CoV-2 Correlate with NET Formation and Fatal Outcome in Severely Diseased COVID-19 Patients. Cells. 2020;9(12).

31. Ma H, Zeng W, He H, Zhao D, Jiang D, Zhou P, et al. Serum IgA, IgM, and IgG responses in COVID-19. Cellular & Molecular Immunology. 2020;17(7):773–5.

32. Padoan A, Sciacovelli L, Basso D, Negrini D, Zuin S, Cosma C, et al. IgA-Ab response to spike glycoprotein of SARS-CoV-2 in patients with COVID-19: A longitudinal study. Clinica Chimica Acta. 2020;507:164–6.

33. Jiang HW, Li Y, Zhang HN, Wang W, Yang X, Qi H, et al. SARS-CoV-2 proteome microarray for global profiling of COVID-19 specific IgG and IgM responses. Nat Commun. 2020;11(1):3581.

34. de Assis RR, Jain A, Nakajima R, Jasinskas A, Felgner J, Obiero JM, et al. Analysis of SARS-CoV-2 antibodies in COVID-19 convalescent blood using a coronavirus antigen microarray. Nature Communications. 2021;12(1):6.

35. Tillett RL, Sevinsky JR, Hartley PD, Kerwin H, Crawford N, Gorzalski A, et al. Genomic evidence for reinfection with SARS-CoV-2: a case study. Lancet Infect Dis. 2021;21(1):52–8.

36. Prado-Vivar B, Becerra-Wong M, Guadalupe JJ, Marquez S, Gutierrez B, Rojas-Silva P, et al. COVID-19 Re-Infection by a Phylogenetically Distinct SARS-CoV-2 Variant, First Confirmed Event in South America. SSRN; 2020.

